# Whole-Genome RNA Sequencing of Femoral Head Impingement Cartilage Identifies FGF18 As A Biomarker in Hip Osteoarthritis Progression

**DOI:** 10.1101/2022.05.23.492951

**Authors:** Benjamin D. Kuhns, John M. Reuter, Victoria L. Hansen, Gillian Soles, Jennifer H. Jonason, Cheryl Ackert-Bicknell, Chia-Lung Wu, Brian D. Giordano

## Abstract

**Introduction:** The natural history of Femoroacetabular Impingement (FAI) has been clinically associated with the development of hip osteoarthritis (OA); however, the pathobiological mechanisms underlying the transition from focal impingement to global joint degeneration remain unclear. The goal of the study was to investigate differences in transcriptomic profiles of the cartilage from FAI and hip OA patients using whole-genome RNA sequencing.

**Methods:** Thirty-seven patients were included in the study with 20 diagnosed with FAI undergoing arthroscopic treatment and 15 diagnosed with hip OA undergoing total hip arthroplasty (THA). Cartilage samples were obtained intraoperatively over the femoral head-neck junction for both FAI and OA cohorts. Whole-genome RNA sequencing was performed on 10 gender-matched patients in the FAI and OA cohorts with the remaining samples used for histopathologic analysis using the Osteoarthritis Research Society International (OARSI) grading system and qRT-PCR validation. Target validation was further confirmed with immunohistochemical staining for FGF18 on FAI and OA cartilage samples.

**Results:** We identified a total of 3,531 Differentially Expressed Genes (DEGs) between the FAI and OA cohorts with multiple targets for genes implicated in canonical OA pathways. qRT-PCR validation confirmed increased expression of FGF18 and WNT16 in the FAI samples, while there was increased expression of MMP13 and ADAMTS4 in the OA samples. Expression levels of FGF18 and WNT16 were also higher in FAI samples with mild cartilage damage (OARSI grades 1-2) compared to FAI samples with severe cartilage damage or OA cartilage (OARSI grade 4-6). Immunohistochemical evaluation identified increased FGF18 staining in OARSI grade 1-2 FAI samples compared to OARSI grade 5-6 FAI and OA samples.

**Conclusions:** RNA sequencing of cartilage of FAI and hip OA patients identified a negative association of FGF18 expression levels with cartilage damage severity, suggesting that FGF18 may be used as a marker for hip OA progression. Future evaluation of FGF18 signaling as well as other markers in early OA may yield further insight into disease prevention and treatment.

## Introduction

Femoroacetabular Impingement (FAI) is a common syndrome causing hip pain and dysfunction in young and middle-aged adults while hip osteoarthritis (OA) typically affects older patients and represents a process of global joint deterioration. The cam deformity that defines FAI presents as an incongruence at the femoral head-neck junction that impinges on the chondrolabral soft tissues with hip flexion and rotation. Recent studies have found a strong correlation between the size of the cam deformity and OA progression.^1–5^ Despite this clinical association, the specific pathophysiologic mechanisms responsible for disease progression are largely unknown, and the transition from a process defined by focal impingement to one of global joint deterioration remains an area of active investigation.

Several studies have previously evaluated the expression of select genes isolated from pathologic tissues in FAI and hip OA (Table 1). Taken together, these findings suggest that there are distinct expression profiles of FAI and OA articular cartilage with regards to mediators of inflammation and cartilage homeostasis. Further insight into altered gene expression between FAI and OA cartilage may identify unique biomarkers associated with OA progression and yield insight into novel treatment strategies for early OA in at-risk hips.^6,7^

**Table 1:** Differential expression for previously identified markers in FAI and OA.^6–12^*indicates increased expression in OA compared to FAI.

| **Cartilage** | **Labrum** | **Synovium** | **Subchondral Bone** | **Serum** |
| --- | --- | --- | --- | --- |
| IL-1β | IL-1β * | IL-1β * | IL-6 | COMP |
| IL-8 | IL-8* | IL-8* | RANKL | CRP |
| CCL3L1 | MMP-3* | MMP-3* | OPG |  |
| MMP13 | COL1A1* |  | ALP |  |
| ADAMTS4 |  |  |  |  |
| ACAN |  |  |  |  |
| COL2A1 |  |  |  |  |
| P21 |  |  |  |  |
| Bcl-2 |  |  |  |  |
| FasL |  |  |  |  |

The purpose of this study was to use whole-genome RNA sequencing to evaluate and validate differentially expressed genes in femoral head articular cartilage in patients with Cam deformities undergoing arthroscopic for FAI compared to patients undergoing total hip arthroplasty (THA) for end stage OA secondary to FAI. We hypothesized that FAI cartilage would have a distinct gene expression profile compared to osteoarthritic cartilage.

## Materials and Methods

### Subject Recruitment

IRB approval was obtained prior to the study with subjects consenting to cartilage sampling preoperatively. Inclusion criteria for the study included a clinical and radiographic diagnosis of femoroacetabular impingement or end-stage OA requiring surgery as well as an alpha angle (AA) of >60°. Exclusion criteria included prior hip surgery, dysplasia (LCEA >25°), avascular necrosis, and rheumatologic conditions. Subject demographics were collected by review of the electronic medical record. AP pelvis radiographs were evaluated to determine AA, lateral center edge angle (LCEA), and Tonnis grade and were performed by the lead and senior authors. Intraoperative cartilage assessment was performed using the Outerbridge classification.^8^

### Cartilage Acquisition

Cartilage samples were harvested intraoperatively from both the FAI and OA cohorts. For FAI samples, the location of the CAM deformity over the anterosuperior femoral head-neck junction was confirmed fluoroscopically. It was then excised with osteotomes and transferred to the back table where any residual subchondral bone was then sharply debrided with a scalpel prior to flash freezing with liquid nitrogen. For OA samples, once the femoral head was excised it was transferred to the back table where a 10mm biopsy punch was used to obtain an osteochondral sample overlying the anterosuperior femoral head-neck junction. Subchondral bone was then sharply excised, and the cartilage specimen was flash frozen in liquid nitrogen. Where possible, additional sample was acquired for histologic analysis and these samples were immediately transferred to 10% neutral buffered formalin for fixation.

### RNA Isolation and Sequencing

Articular cartilage RNA isolation was performed as previously described by Le Bleu et al. (Appendix 1). Briefly, cartilage samples were cryogenically pulverized followed by RNA isolation with TRIzol reagent (Invitrogen) with lysate purified through GeneJET RNA purification kit (ThermoFisher). RNA integrity number scores as well as average A_260/280_ and A_260/230_ ratios were used to determine RNA quality and evaluate fitness for downstream analysis. RNA sequencing was then performed in conjunction with the University of Rochester Genomic Research Center (Appendix 2).

### Sequencing Validation

Isolated RNA was transcribed into cDNA. Gene expression was then quantified by using real-time qPCR reactions performed with using the QuantiTect SYBR Green PCR kit (Qiagen; Hilden, Germany). We used gene specific primers (Supplemental Table S1; Integrated DNA Technologies Inc; Coralville, Iowa). Transcript quantity measurements were normalized to GADPH and gene expression levels were quantified using the 2^^(-ΔΔCT)^ method.

### Histology Immunohistochemistry

Following fixation for 48-72 hours, specimens were dehydrated and embedded in paraffin wax followed by sectioning. Sections were deparaffinized and stained for Hematoxylin & Eosin, Safranin-O, and Anti-FGF18 respectively Appendix 3).

Histologic cartilage quality was graded in a blinded fashion by 2 faculty investigators (CLW, JJ) using the OARSI criteria which has previously demonstrated satisfactory intra and inter-observer reliability. The intraclass correlation coefficient (ICC) between the faculty graders was 0.74 with grading differences resolved in conference for the final analysis. FAI samples that had lower OARSI scores (1-2) were considered low-grade FAI while FAI samples with higher OARSI scores (4-5) were considered high-grade FAI. A post-sequencing validation cohort of 4 low-grade, and 4 high-grade OARSI cartilage samples from FAI patients was compared to 4 OA cartilage samples with qRT-PCR evaluation of select markers (Supplemental Table S1).

### Statistical Analysis

The results of qRT-PCR and histological grading were analyzed with JMP (Cary, NC) and PRISM (San Diego, CA). Differential expression was determined using the false discovery rate (FDR) or adjusted p value < 0.05, as appropriate. A priori power analysis (G*Power; Universität Düsseldorf^9^) was performed using the aligned data from the RNA sequencing results on FGF 18 which required 3 subjects in the FAI and OA cohorts to determine differential expression with adequate power (1-β>0.8; p<0.05).

## Results

### Cohort Demographics

37 subjects (15 female; 40%) with a mean age of 46.5±17.5 and BMI of 28.7±5.9 were included in the study (Table 1). Cartilage samples from 20 subjects yielded high quality RNA suitable for downstream analysis and were included in the RNA sequencing arm of the study. Cartilage samples harvested from the remaining 17 subjects were used for RNA sequencing validation with qRT-PCR and IHC analyses. Subjects in the OA cohort were significantly older with a higher BMI however there was no difference in AA or LCEA (Table 2).

**Table 2:** Cohort Demographics. FAI: Femoroacetabular Impingement; OA: Osteoarthritis; BMI: Body Mass Index; AA: Alpha Angle; LCEA: Lateral Center Edge Angle

|  | **FAI** | **OA** | **p** |
| --- | --- | --- | --- |
| *Study Cohort* |  |  |  |
| **n** | 21 | 16 |  |
| **Gender (F)** | 8 (38%) | 7 (44%) | 0.75 |
| **Age** | 34.2±11.5 | 62.7±8.3 | <0.001 |
| **BMI** | 26.6±4.7 | 31.5±6.2 | 0.01 |
| **AA** | 66.7±5.3 | 65.4±3.6 | 0.41 |
| **LCEA** | 36.2±6.0 | 33.7±5.1 | 0.2 |
| *Sequencing Cohort* |  |  |  |
| **n** | 10 | 10 |  |
| **Gender (F)** | 5 (50%) | 5 (50%) | 1 |
| **Age** | 34.0±13.4 | 63.1±8.0 | <0.001 |
| **BMI** | 25.8±4.8 | 31.3±6.3 | 0.04 |
| **Alpha Angle** | 66.0±3.5 | 65.1±3.9 | 0.61 |
| **LCEA** | 35.9±5.8 | 33.0±4.1 | 0.22 |
| *Validation Cohort* |  |  |  |
| **n** | 11 | 6 |  |
| **Gender (F)** | 3 (27%) | 2 (33%) | 0.79 |
| **Age** | 34.4±10.1 | 62.0±9.6 | <0.001 |
| **BMI** | 27.2±4.7 | 31.8±6.6 | 0.11 |
| **Alpha Angle** | 67.3±6.6 | 65.9±3.5 | 0.63 |
| **LCEA** | 36.4±6.5 | 34.9±6.7 | 0.66 |

All patients in the FAI cohort had Tönnis grades 0-1 while the OA cohort had predominantly grade 3 degenerative changes (Table 3). The OARSI grades for FAI ranged from 1-5 while the OARSI grades for OA cartilage ranged from 2-6. Arthroscopic cartilage evaluation of FAI patients identified grades 0-IV degeneration on both the acetabulum and femoral head. There were no significant differences between individual OARSI grades for the FAI and OA cohorts (p>0.05). Low-grade FAI impingement cartilage samples with mild degenerative changes (OARSI grade 1-2) were compared to high grade FAI samples with severe cartilage degeneration (OARSI grade 4-5) as well as osteoarthritic cartilage samples in a validation cohort (OARSI grade 4-5; Table 4). Low-grade cartilage specimens had a greater proportion of arthroscopic femoral head Outerbridge Grade 0 changes (100%) compared to high grade FAI samples (0%; p=0.008; Supplemental table S2) and there was a significant association between increasing histologic OARSI grade and intraoperative femoral head cartilage grade (r=0.68; p<0.01). There were no differences in Outerbridge acetabular cartilage grading when comparing low-grade and high-grade FAI specimens.

**Table 3:** Radiographic, Intraoperative and immunohistochemical cartilage assessment. FAI: Femoroacetabular Impingement. OA: Osteoarthritis. OARSI: Osteoarthritis Research Society International

|  | **FAI** | **OA** |
| --- | --- | --- |
| **Tonnis Grade** |  |  |
| **0** | 12 (58%) | 0 |
| **1** | 9 (42%) | 0 |
| **2** | 0 | 4 (25%) |
| **3** | 0 | 12 (75%) |
| **OARSI Cartilage Grade** |  |  |
| **1** | 1 (6.5%) | 0 (0%) |
| **2** | 5 (33%) | 1 (10%) |
| **3** | 2 (13%) | 1 (10%) |
| **4** | 3 (20%) | 3 (30%) |
| **5** | 4 (26.5%) | 4 (40%) |
| **6** | 0 (0%) | 1 (10%) |
| **Outerbridge Femoral Head Cartilage Grade** |  |  |
| **0** | 9 (43%) |  |
| **1** | 4 (19%) |  |
| **2** | 2 (9.5%) |  |
| **3** | 2 (9.5%) |  |
| **4** | 4 (19%) |  |
| **Outerbridge Acetabular Cartilage Grade** |  |  |
| **0** | 0 (0%) |  |
| **1** | 2 (9.5%) |  |
| **2** | 12 (57%) |  |
| **3** | 3 (14.5%) |  |
| **4** | 4 (19%) |  |

**Table 4.** Demographic comparison of Low-Grade FAI, High Grade-FAI, and OA patients in the qRT-PCR validation cohort. FAI: Femoraacetabular impingement; OA: Osteoarthritis; BMI: Body Mass Index; AA: Alpha Angle; LCEA: Lateral Center Edge Angle p* Low grade FAI compared to high-grade FAI. P** Low grade FAI compared to OA p*** High Grade FAI compared to OA

|  | Low-Grade FAI | High-Grade FAI | OA | P* | p** | p*** |
| --- | --- | --- | --- | --- | --- | --- |
| **n** | 4 | 4 | 4 |  |  |  |
| **Gender (F)** | 2 (50%) | 1 (25%) | 2 (50%) | 0.49 | 1.0 | 0.49 |
| **Age** | 31.7±3.6 | 40.2±14.8 | 65.3±2.9 | 0.92 | 0.001 | 0.001 |
| **BMI** | 25.1±6.5 | 29.4±3.8 | 32.2±8.1 | 0.58 | 0.39 | 0.19 |
| **AA** | 65.6±3.1 | 66.6±10.3 | 66.8±3.9 | 0.26 | 0.62 | 0.37 |
| **LCEA** | 33.1±7.3 | 37.4±7.9 | 36.7±7.3 | 0.52 | 0.94 | 0.61 |

### RNA Sequencing and Validation

Whole-genome RNA sequencing identified 3,532 genes that were significantly differentially expressed between the FAI and OA sequencing cohorts (p < 0.05; Figure 1). Of these, there were 27 genes in the OA cohort that had a Log_2_ fold change (FC) greater than 2 while there were 523 genes that had a Log_2_FC greater than 2. When the differential expression data was analyzed with the IPA platform, multiple genes were identified that were involved in the canonical OA signaling pathway (Table 5).

**Table 5:** Differential expression of select gene in the canonical osteoarthritis pathway

| Gene | Function and Relation to OA | padj | FC |
| --- | --- | --- | --- |
| *FAI Upregulated* |  |  |  |
| **SMAD4** | Mediates TGFβ signaling<sup>18</sup> | 2.19E-03 | 0.181 |
| **C/EBPβ** | Transcription factor, increases Type X Collagen expression, increased in early OA<sup>19</sup> | 2.32E-04 | 0.399 |
| **WNT16** | Wnt Antagonist, Chondroprotective<sup>20</sup> | 1.93E-03 | 1.528 |
| **FGF18** | Anabolic growth factor chondroprotective against OA<sup>21,22</sup> | 9.67E-09 | 2.038 |
| **NOS1** | Pro-inflammatory, increase increased in OA<sup>23</sup> | 1.87E-34 | 4.898 |
| **GDF6** | Ligand augmenting BMP signaling<sup>24</sup> | 1.08E-07 | 2.567 |
| **BMPR1B** | BMP receptor, chondroprotective<sup>25</sup> | 1.31E-22 | 2.036 |
| **HIF3A** | Transcription factor inhibiting cartilage catabolism<sup>26</sup> | 1.20E-08 | 1.901 |
| *OA upregulated* |  |  |  |
| **GREM1** | BMP antagonist, chondroprotective<sup>27</sup> | 3.32E-12 | -3.291 |
| **RANK** | Increased in response to IL-1β signaling in OA<sup>28</sup> | 7.70E-08 | -1.709 |
| **COL10A1** | Short chain collagen diffusely increased in OA<sup>29</sup> | 1.64E-04 | -2.222 |
| **MMP13** | Cleaves type 2 collagen, pro-inflammatory, increased in OA<sup>30</sup> | 2.77E-04 | -1.935 |
| **ADAMTS4** | Enzyme degrading aggrecan in cartilage ECM, pro-inflammatory, increased in OA<sup>31</sup> | 7.35E-03 | -1.376 |
| **TNF** | Pro-Inflammatory cytokine, increased in OA<sup>32</sup> | 6.12E-03 | -1.129 |
| **RUNX2** | Transcription Factor, increased in OA<sup>33</sup> | 4.76E-03 | -1.114 |
| **TGFB1** | Induces osteophyte formation and angiogenesis, increased in OA<sup>34</sup> | 1.38E-03 | -0.461 |

**Figure 1:**
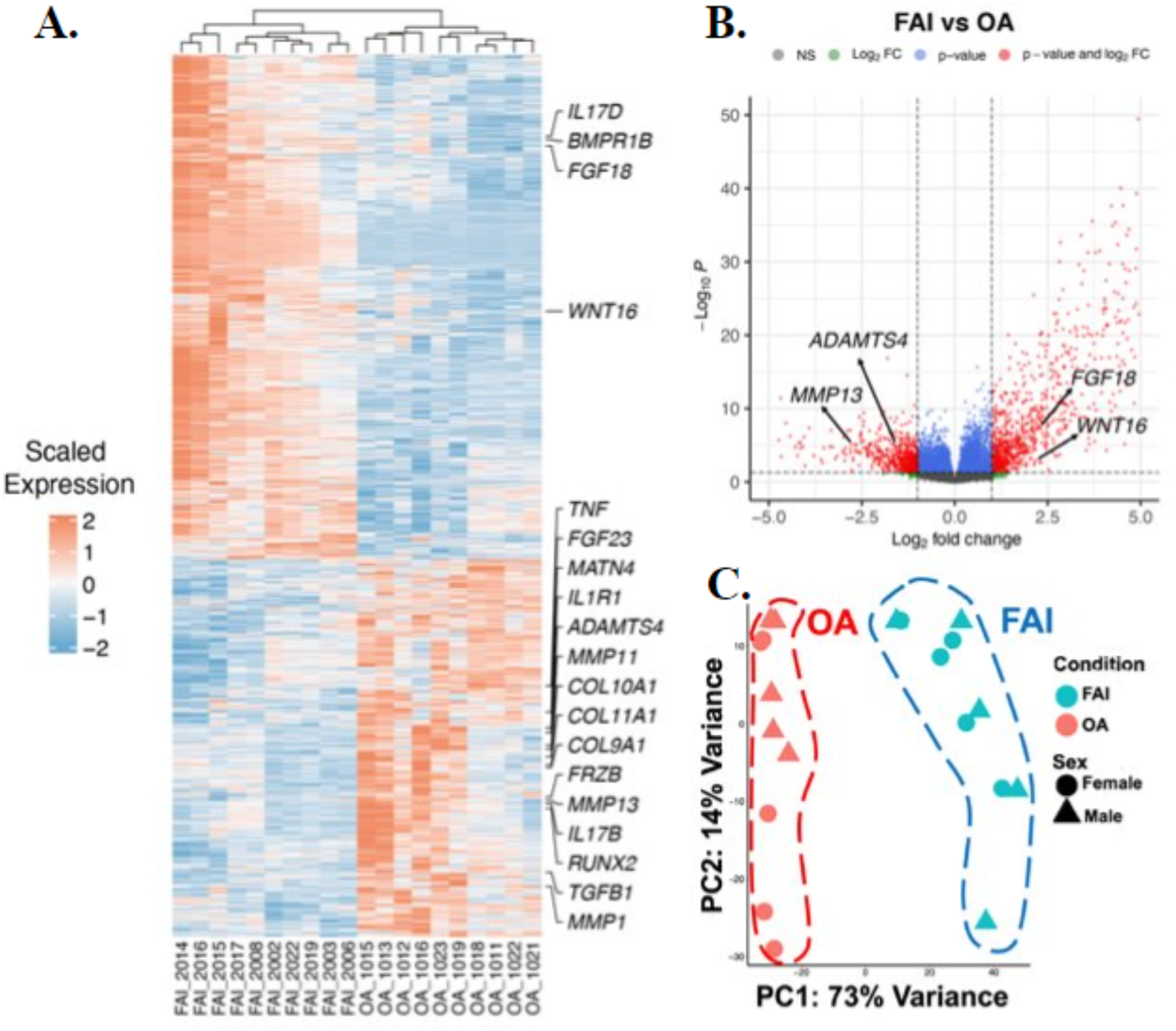
A: Heat map demonstrating sample specific differential expression (FAI, red, positive log fold change), OA (Blue, negative log fold change). B. Volcano plot demonstrating differential gene expression between FAI and OA cohorts. Negative fold change represents increased expression of genes in the OA cohort while positive fold change represents increased expression of genes in the FAI cohort. C. Primary component analysis (PCA) demonstrating sequencing variation between OA and FAI samples. NS: Not significant. Log2FC: Log_2_ Fold Change highlighting p-value: FDR >0.05).

### qRT-PCR Target Validation

qRT-PCR analysis was performed on select biomarkers (FGF18; WNT16; MMP13; ADAMTS4) from low-grade and high-grade FAI impingement cartilage as well as the OA cartilage samples in the validation cohort. Low-grade FAI cartilage demonstrated a 343.1-fold increase in expression compared to OA cartilage, while high-grade FAI had an 11.0-fold increase in expression compared to OA cartilage (Figure 2A). WNT16 was increased 57.8-fold in low-grade FAI and 19.0-fold in high-grade FAI with respect to OA cartilage (Figure 2B). OA cartilage had significantly increased expression of MMP13 compared to early and late FAI cartilage and significantly increased expression of ADAMTS-4 over early FAI cartilage (Figure 2C-D; p<0.05). MMP13 also had significantly greater expression in late, compared to early FAI (p<0.05).

**Figure 2:**
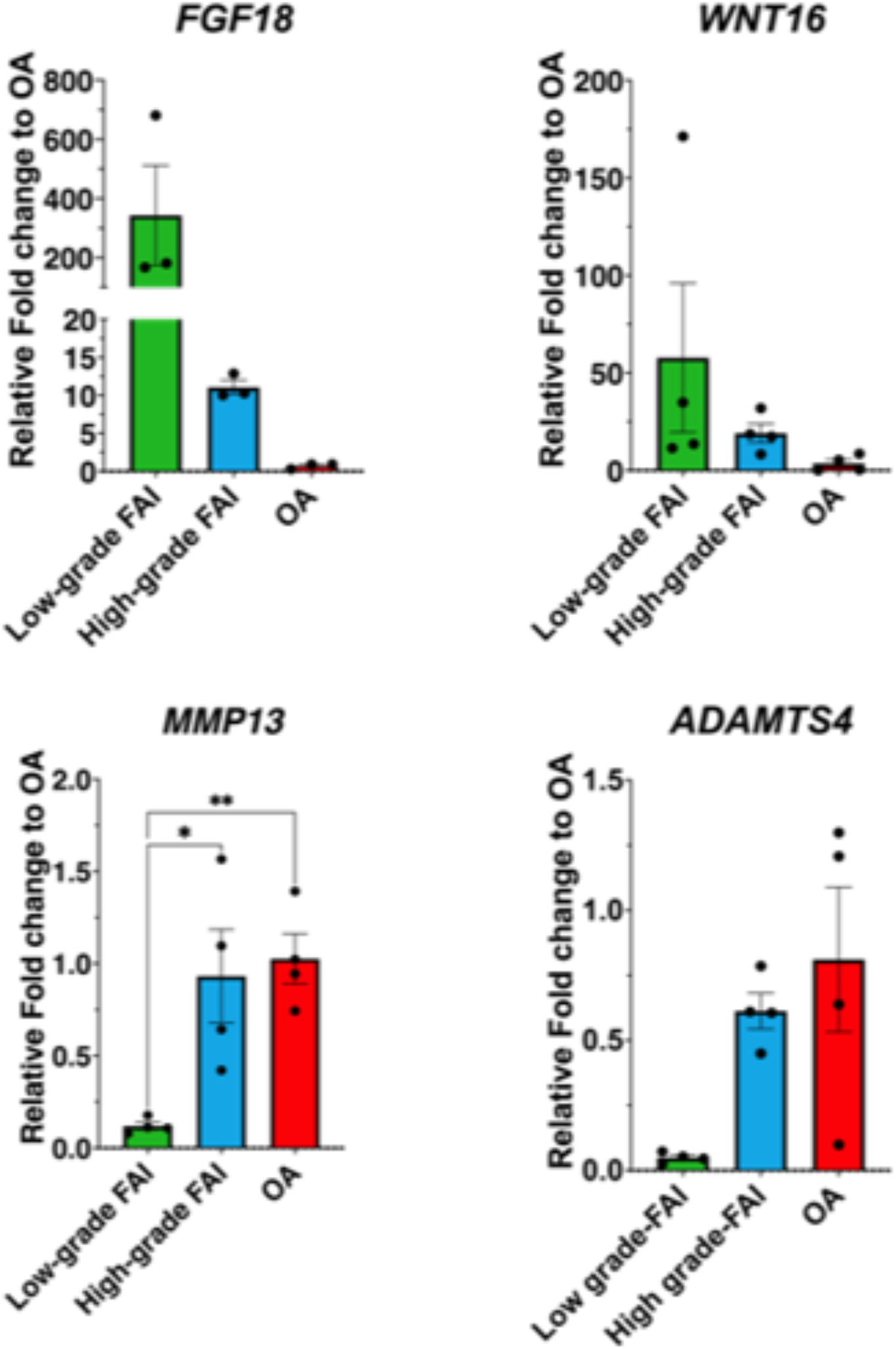
qRT-PCR validation for specific markers identified in RNA sequencing. Cartilage expression of FGF18 and WNT16 were upregulated in FAI, particularly low-grade FAI, while MMP13 and ADAMTS4 were upregulated in osteoarthritic and high-grade FAI cartilage samples.

### Immunohistochemical target confirmation

Immunohistochemical staining against FGF18 was performed on articular cartilage sections from the low-grade and high-grade FAI impingement cartilage as well as OA samples in the validation cohort. FGF18 expression was increased in patients with low grade cartilage degeneration compared to FAI patients with high grade cartilage lesions (Figure 3).

**Figure 3:**
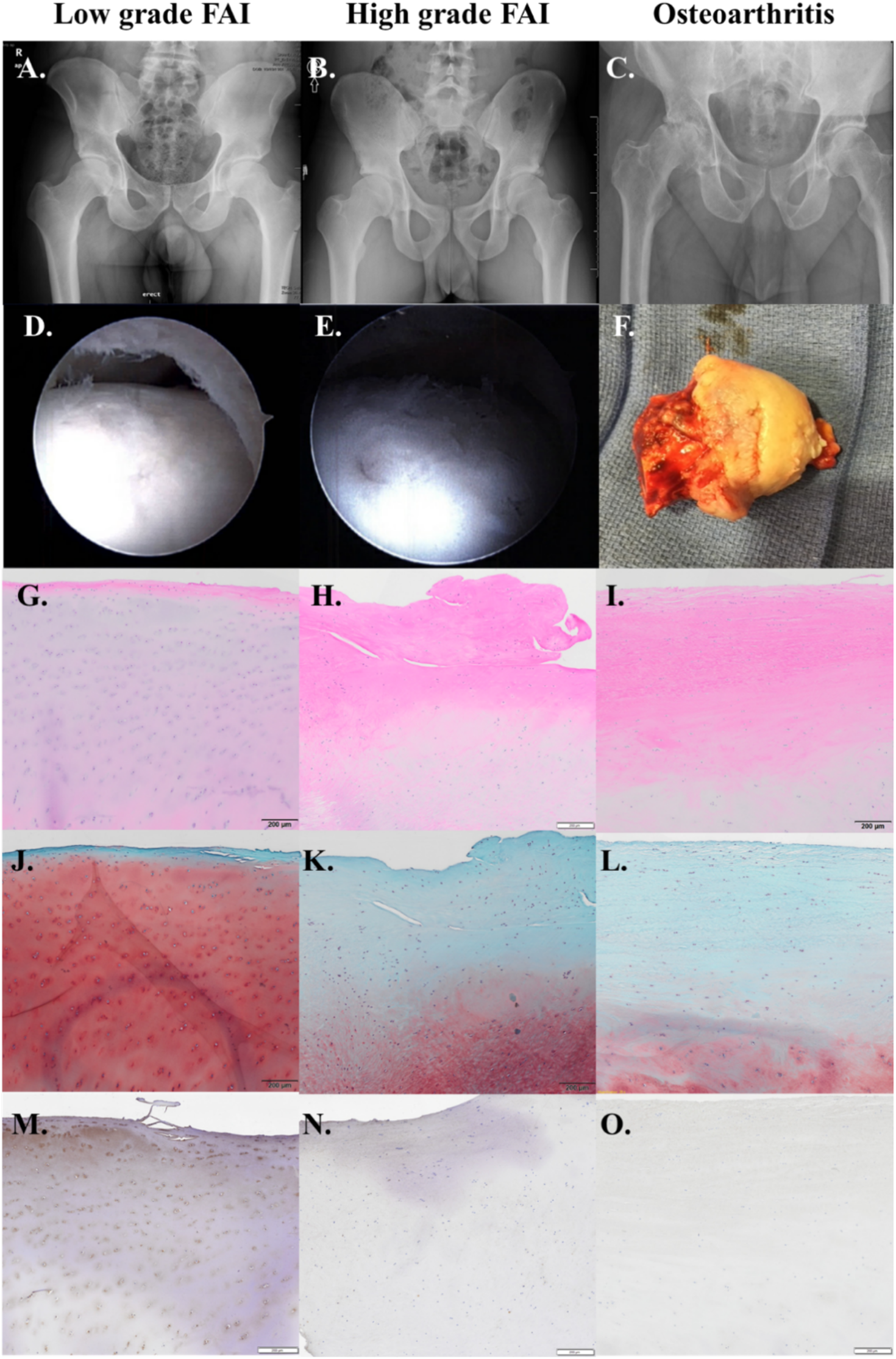
Radiographic (A-C), Intraoperative (D-F), H&E (G-I), Safranin-O (J-L), and anti-FGF 18 immunostaining (M-O) for patients with low grade FAI (OARSI 1), high grade FAI (OARSI 4) and OA (OARSI 5). FGF18 IHC staining was increased in cartilage samples from low grade FAI compared to high grade and osteoarthritic samples. Histologic samples were taken at 5x magnification, scale bar = 200 µm)

## Discussion

The hypothesis that FAI and OA femoral head articular cartilage will have distinct genomic expression profiles was confirmed through whole-genome RNA sequencing of femoral head articular cartilage samples. We identified differential expression of multiple genes that are involved in the canonical OA signaling pathway. We then validated expression data for several markers relevant to OA pathogenesis and progression. We found that FAI cartilage with lower OARSI grades had elevated FGF18 and WNT16 expression and lower MMP13 and ADAMTS4 expression than FAI cartilage and OA cartilage with higher OARSI grades. We also found that histologic OARSI grades for femoral head cartilage were correlated with intraoperative Outerbridge femoral head cartilage grading but not acetabular cartilage grading.

Differences in inflammation and cartilage metabolism between FAI and OA have been previously reported. In a landmark study by Hashimoto and colleagues, the authors demonstrated increased ADAMTS4, IL-8, and ACAN in FAI patients compared to OA patients that had preoperative FAI morphology but no differences in MMP13 or IL-1β expression.^6^ Further, using Beck’s macroscopic cartilage grading system the authors found increased relative expression of IL-8, ACAN, and COL2A1 in FAI patients with cartilage cleavage compared to distinct cartilage disease states in FAI (chondromalacia; chondral defect or normal appearing) as well as osteoarthritic patients, suggesting that gene expression is altered based on the cartilage status at the time of acquisition.^6^ In a subsequent study by Chinzei et al. the authors found increased expression of MMP13, IL-1β, IL-8, and ADAMTS4 in impingement cartilage in an FAI cohort compared to OA.^10^ The discrepancies in MMP13 and IL-1β expression levels between these studies may possibly result from etiologic differences in the OA cohorts. Hashimoto and colleagues sampled osteoarthritic cartilage in patients with FAI morphology and found no differences in MMP13 and IL-1β expression while Chinzei et al. report significantly increased MMP13 and IL-1β expression in FAI cartilage compared to an osteoarthritic cohort of undefined etiology.^6,10^ It is possible that OA secondary to FAI may have an increased inflammatory profile relative to OA secondary to dysplasia or other etiologies. This is supported by recent immunohistologic data from Haneda and colleagues who found no differences in staining of IL-1β, ADAMTS4, and MMP13 in cartilage samples from patients with FAI or OA secondary to FAI, but both were increased compared to osteoarthritic cartilage obtained from dysplastic hips.^11^ Consistent with these studies, we found that markers of cartilage breakdown (MMP13, ADAMTS-4) were expressed in both FAI and FAI-associated OA cartilage tissues. While we observed increased expression of these markers in the OA compared to the total FAI cohort on the sequencing arm of this study, subsequent validation with qRT-PCR found similar expression between high grade FAI and osteoarthritic cartilage, suggesting that expression of inflammatory markers increases as FAI progresses towards OA. These findings were supported in a recent study by Liang and colleagues who found differences in COL2A1, ACAN, and MMP3 immunostaining between early and late-stage FAI samples.^12^ Taken together, these results suggest that there are both etiologic and temporal factors impacting gene expression between FAI and osteoarthritic cartilage, and as the degenerative processes of FAI become more advanced the articular cartilage expression profile parallels that of osteoarthritic cartilage.

Differences in expression between low OARSI grade FAI and high OARSI grade demonstrate temporal changes associated with advancing cartilage degradation. Impingement cartilage obtained from the FAI population in this study reflects a spectrum of chondral pathology confirmed with macroscopic intraoperative grading (Outerbridge grades 0-4) as well as microscopic immunohistologic evaluation (OARSI grade 0-5). We found potential anabolic and chondroprotective genes (FGF18; WNT16) had greater expression in low grade FAI compared to high grade FAI and OA while catabolic genes (MMP13 and ADAMTS4) had greater expression in high grade FAI and OA compared to low grade FAI. These results are consistent with the recent studies reported by Haneda and colleagues that found FAI cartilage samples with higher OARSI grades to have similar expression of MMP13 and ADAMTS-4, both of which were elevated compared to control samples without OA.^11^ While impingement cartilage studied in that report had similar OARSI grades compared to osteoarthritic samples (4.3 vs. 3.6), we identified several patients with minimal cartilage degradation (OARSI grades 1-2) at the time of acquisition. Understanding the evolution of cartilage gene expression as FAI progresses towards end-stage OA may yield insight into the molecular processes involving the transition from focal impingement to global joint destruction.

A recent genome-wide analyses (GWAS) study identified ten OA associated genes whose encoded proteins have targeted therapeutics in development or already on the market.^13^ RNA sequencing results from this study found significantly differential expression in four of these ten genes (TGFβ1, FGF18, CTSK, MAPT). For this study, we focused on validating FGF18 expression results as FGF18 has been shown to have significant chondrogenic effects in both in vitro and in vivo disease models of OA.^14–21^ Acting through FGFR3 signaling, FGF18 has been shown to increase type II collagen production as well as stimulate cartilage repair, increase cartilage thickness, and prevent joint degradation in murine models of post-traumatic OA.^14,16,17,21^ Further, Sprifermin (the recombinant human FGF18 analog drug) has been shown to promote a dose dependent increase in cartilage thickness, particularly in the lateral compartment, of patients with pre-existing knee OA in the double-blind randomized control FORWARD (FGF18 Osteoarthritis Randomized Trial with Administration of Repeated Doses) drug trial.^19^ Eckstein and colleagues recently reported follow up data from the FORWARD trial where they found that patients at higher risk for OA progression receiving 100 µg of Sprifermin every 6 months had maintained cartilage thickness and decreased WOMAC scores compared to the placebo group at 5 year follow-up.^22^ Additionally, no patients in the 100µg per 6 month group required a knee replacement compared to 4.6% in the placebo group at the 5 year time point. The finding from the present study that FGF18 is upregulated in FAI, particularly low-grade FAI, compared to OA cartilage indicates that altered FGF18 signaling may play a role in the progression of FAI induced OA. Additional studies are required to further evaluate FGF18 signaling in FAI to investigate downstream signaling effects, additional therapeutic targets, and the role of Sprifermin in hip cartilage repair and OA prevention.

### Limitations

This study has multiple limitations that may affect the interpretation of its findings. There was not a true negative control cohort of patients without FAI or OA, as harvesting cartilage from asymptomatic donors without evidence of disease would be unethical. Second, while efforts were made to ensure that patients in the OA cohort had Cam morphology, we were unable to obtain radiographs prior to OA onset and it is possible that the increased AA was secondary to osteophyte formation. There were also differences in sampling techniques, as tissues from FAI patients were collected arthroscopically while tissues from OA patients were harvested after femoral head resection during total hip arthroplasty. Additionally, while both PCR and IHC specimens in the validation cohort were obtained from the same region of the Cam deformity at the anterolateral femoral head-neck junction, it is possible that there were geographic expression differences within the Cam deformity itself. Future studies will be required to evaluate the homogeneity of gene expression and cartilage quality throughout the Cam deformity. Despite these limitations, using an unbiased whole-genome sequencing technique, the present study identified significantly differential expression between FAI and OA in multiple signaling pathways relevant to OA.

## Conclusion

Our RNA sequencing of articular cartilage in the impingement zone of FAI and OA patients revealed significantly differential expression of greater than 3000 genes. Our results also uncovered distinct gene expression profiles and signaling pathways between FAI and OA cartilage that may reflect hip OA progression. A detailed understanding of molecular mechanisms underlying transcriptomic reprograming in chondrocytes from FAI to OA will provide significant insight into the development of early therapeutic intervention for hip OA.

