## Supplemental files for "Whole-Genome RNA Sequencing of Femoral Head Impingement Cartilage Identifies FGF18 As A Biomarker in Hip Osteoarthritis Progression"

| Primer | Sequences |
| --- | --- |
| GAPDH | 5'-CCACATCGCTGAGACACCAT-3' |
| | 5'-GCGCCCAATACGACCAAATC-3' |
| ADAMTS4 | 5'-GATCCACAACCATGGTGCCT-3' |
| | 5'-CAACCAAAGGAACGGACCCT-3' |
| WNT16 | 5'-TACAGCTCCCTGCAAACGAG-3' |
| | 5'-AGCACCAAGTTATCCCTCGC-3' |
| FGF18 | 5'-GTACGTGGGCTTCACCAAGA-3' |
| | 5'-TGGTCACCGTCGTGTACTTG-3' |
| MMP13 | 5’-TCAAGATGCATCCAGGGGTC-3' |
| | 5’-TCTCGGAGCCTCTCAGTCAT-3' |

**Supplemental Table S1:** Primer sequencing used for RT-PCR.

| OARSI Grade | Low-Grade FAI | High-Grade FAI | OA |
|---|---|---|---|
| **1** | 1 (25%) | 0 (0%) | 0 (0%) |
| **2** | 3 (75%) | 0 (0%) | 0 (0%) |
| **3** | 0 (0%) | 0 (0%) | 0 (0%) |
| **4** | 0 (0%) | 1 (25%) | 1 (25%) |
| **5** | 0 (0%) | 3 (75%) | 2 (50%) |
| **6** | 0 (0%) | 0 (0%) | 1 (25%) |
| **Femoral Head Cartilage*** | | | |
| **0** | 3 (75%) | 0 (0%) | |
| **1** | 1 (25%) | 0 (0%) | |
| **2** | 0 (0%) | 1 (25%) | |
| **3** | 0 (0%) | 0 (0%) | |
| **4** | 0 (0%) | 3(75%) | |
| **Acetabular Cartilage*** | | | |
| **0** | 0 (0%) | 0 (0%) | |
| **1** | 1 (25%) | 0 (0%) | |
| **2** | 2 (50%) | 2 (50%) | |
| **3** | 0 (0%) | 1 (25%) | |
| **4** | 1 (25% | 1 (25%) | |

**Supplemental Table 2:** OARSI histologic grading and Outerbridge arthroscopic cartilage evaluation on a validation cohort of low-grade FAI, high-grade FAI, and OA femoral head cartilage samples. FAI: Femoroacetabular impingement; OA: Osteoarthritis.

**Appendix 1: RNA Isolation**

Harvested articular cartilage was transferred from a liquid nitrogen container after harvesting and pulverized using a cryogenic mill (SPEX SamplePrep 6870 Freezer/Mill®). Extreme care was taken to minimize tissue exposure to the ambient environment. The optimized grinding parameterse included a total of four 3-min cycles of grinding, each at a frequency of 15 cycles per second (cps) followed by 2 min of cool down. One hundred mg of pulverized tissue was then transferred directly to an Eppendorf tube containing 1.5 ml TRIzol and incubated for 15 min at 4°C on an end-to-end rotator. Samples were then centrifuged at 12,000 × g for 5 min at 4°C to pellet undigested tissue. Supernatant was then transferred to fresh Eppendorf tubes and mixed with 200 µL of chloroform by vigorous shaking for 30 seconds, left to stand for 3 min at room temperature, and then centrifuged at 12,000 × g for 15 min at 4°C. The aqueous layer was transferred to a new tube and a mixture of concentrated sodium chloride and sodium acetate was added to achieve final concentrations of 1.2 M and 0.8 M, respectively. This was then purified through GeneJET RNA purification kit (ThermoFisher) according to the manufacturer's specifications.

**Appendix 2: Whole Genome RNA Sequencing**

Total RNA was isolated using the RNeasy Plus Micro Kit (Qiagen, Valencia, CA) per manufacturer's recommendations. RNA concentration was determined with the NanopDrop 1000 spectrophotometer (NanoDrop, Wilmington, DE) and RNA quality assessed with the Agilent Bioanalyzer 2100 (Agilent, Santa Clara, CA). The TruSeq Stranded mRNA Sample Preparation Kit (Illumina, San Diego, CA) was used for next generation sequencing library construction per manufacturer's protocols. Briefly, mRNA was purified from 200ng total RNA with oligo-dT magnetic beads and fragmented. First-strand cDNA synthesis was performed with random hexamer priming followed by second-strand cDNA synthesis using dUTP incorporation for strand marking. End repair and 3` adenylation was then performed on the double stranded cDNA. Illumina adaptors were ligated to both ends of the cDNA, purified by gel electrophoresis and amplified with PCR primers specific to the adaptor sequences to generate cDNA amplicons of approximately 200-500bp in size. The amplified libraries were hybridized to the Illumina flow cell and paired end reads of 101nt were generated for each sample using Illumina's NovaSeq 6000 sequencer. Raw reads generated from the Illumina base calls were demultiplexed using bcl2fastq version 2.19.0. Quality filtering and adapter removal are performed using Trimmomatic-0.36 with the following parameters: "TRAILING:13 LEADING:13 ILLUMINACLIP:adapters.fasta:2:30:10 SLIDINGWINDOW:4:20 MINLEN:35".

Processed/cleaned reads were then mapped to the Homo sapiens reference genome (GRCh38 + Gencode-28 Annotation) using STAR_2.7.0f with the following parameters: --twopassMode Basic --runMode alignReads --genomeDir GENOME --readFilesIn SAMPLE --outSAMtype BAM".[14] Gene level read quantification was derived using the subread-1.6.1 package (featureCounts) with a GTF annotation file (Gencode-28) and the following parameters: "-s 2 -t exon -g gene_name". Differential expression analysis was performed using DESeq2-1.16.1 with a P-value threshold of 0.05 within R version 3.4.1 (https://www.R-project.org/).[15] A PCA plot was created within R using the pcaExplorer to measure sample expression variance. Heatmaps were generated using the pheatmap package were given the rLog transformed expression values. Gene ontology analyses were performed using the EnrichR package.[16] The Ingenuity Pathway Analysis (Qiagen; https://www.qiagenbioinformatics.com/products/ingenuitypathway-analysis) and supplemental literature review were used to correlate the differential expression data to the existing canonical OA signaling pathways

**Appendix 3: Histologic and Immunohistologic Staining**

Sections were deparaffinized in 3 changes of xylenes, and then rehydrated in 2 changes of 100% ethanol, 2 changes of 95% ethanol, and 70% ethanol for 5 min each. H&E slides were rinsed in distilled water and then stained with Mayers hematoxylin for 1 min before being rinsed in tap water until it ran clear. Then the H&E slides were rinsed in 1x PBS, 3 changes of distilled water, and counterstained with alcoholic eosin solution for 1 min. For Safranin-O slides were rinsed in tap water and then stained with Harris's hematoxylin for 3 min and rinsed in tap water until it ran clear. Then Safranin-O slides were treated with acid alcohol Differentiation Solution (Sigma, St Louis, MO) for 15 sec and rinsed in tap water for 3 min. Then Safranin-O slides were stained with 0.02% aqueous Fast Green FCF (Electron Microscopy Sciences, Hatfield, PA) solution for 5 min, rinsed briefly in 1% glacial acetic acid, and rinsed in tap water. Safranin-O slides were stained with 0.1% aqueous Safranin O Solution (Sigma-Aldrich, Hanover, Germany) for 3 min and rinsed in 2 changes of 100% ethanol. Anti-FGF18 immunohistochemistry slides were rinsed in deionized water (diH2O) and antigen retrieval was performed using Pepsin Reagent (Sigma, St Louis, MO) incubated for 5 min at room temperature (RT). Slides were rinsed in diH2O and endogenous peroxidases were blocked using  a 1:10 dilution of 30% hydrogen peroxide (Sigma, St Louis, MO) in methanol solution for 10 min at RT. Slides were rinsed in 3 changes of 1x PBS and non-specific antibody binding was blocked using 10% normal goat serum (EMD Millipore, Temecula, CA) for 1 hour at RT. Rabbit anti-human FGF18 antibody (SAB4503479, Sigma, St Louis, MO, 1:50 dilution in 2.5% normal goat serum) was added to each section and incubated overnight at 4°C. Negative control slides were instead incubated with just 2.5% normal goat serum (Vector Laboratories, Burlingame, CA) overnight at 4°C. The next day slides were rinsed in 3 changes of 1x PBS and goat anti-rabbit IgG (Invitrogen ,Rockford, IL, 1:8000 dilution in 2.5% normal goat serum) was added to all sections and incubated for 2 hours at RT. The slides were rinsed with 3 changes of 1x PBS and Vectastain Elite ABC for proxidase solution (Vector Laboratories, Burlingame, CA) was added to each slide and incubated for 30 min at RT. The slides were rinsed with 3 changes of 1x PBS and 2 changes of diH2O. ImmPACT DAB Substrate for peroxidase (Vector Laboratories, Burlingame, CA)  was added to each slide and incubated for 10 min at RT while monitoring the color change and then the reaction was quenched in diH2O. Slides were counter-stained with Hematoxylin QS (Vector Laboratories, Burlingame, CA). H&E, Safaranin-O, and anti-FGF18 slides were all dehydrated in 70% ethanol, 2 changes of 95% ethanol, 2 changes of 100% ethanol, and 3 changes of xylenes for 2 min each and coverslipped with Permount (Fisher Chemical, Fair Lawn, NJ). All microscope images were taken using a V120 Slide Scanner (Olympus) and VS-ASW imaging software (v2.9.2, Olympus).
